# Bioactivity assessment of natural compounds using machine learning models based on drug target similarity

**DOI:** 10.1101/2020.11.06.371112

**Authors:** Vinita Periwal, Stefan Bassler, Sergej Andrejev, Natalia Gabrielli, Kaustubh Raosaheb Patil, Athanasios Typas, Kiran Raosaheb Patil

## Abstract

Natural compounds constitute a rich resource of potential small-molecule therapeutics. While experimental access to this resource is limited due to its vast diversity and difficulties in systematic purification, computational assessment of structural similarity with known therapeutic molecules offers a scalable approach. Here, we assessed functional similarity between natural compounds and approved drugs by combining multiple chemical similarity metrics and physicochemical properties through a random forest model. As a training set, we used pair-wise similarity between 1410 drugs in terms of their shared protein targets. The resulting model featured high performance metrics (matthews correlation coefficient of 0.81, and balanced accuracy of 0.91) suggesting that it well-captured the structure-activity relation. The model was then used to predict protein targets of circa 11k natural compounds by comparing them with the drugs. This revealed therapeutic potential of several natural compounds, including those with support from previously published sources as well as those hitherto unexplored. We experimentally validated one of the predicted link’s activities, viz., Cox-1 inhibition by 5-methoxysalicylic acid, a molecule commonly found in tea, herbs and spices. In contrast, another natural compound, 4-isopropylbenzoic acid, which showed a higher similarity when considering the most weighted similarity metric but was not picked by the random forest model, did not inhibit Cox-1. Our results demonstrate the utility of a machine-learning approach combining multiple chemical features for uncovering protein binding potential of natural compounds.

## Introduction

Around 65% of the small-molecule drugs in use today have originated from natural compounds or their derivatives [1]. Therapeutic effects of natural compounds are thus central to drug discovery [2-5]. Further, identification of bioactive compounds present in the diet and their effect on health has been an active area of research since long [6, 7]. A number of recent studies have reported that dietary natural compounds (such as polyphenols, alkaloids) can reduce the risk of many chronic diseases [8-10], lead to drug and food interactions [11-13], and significantly alter or diversify the composition of the human gut microbiome [14-17]. While natural compounds possess rich structural diversity, often have selective biological actions, and are prevalidated on various biological targets by evolutionary selection [18-21], they are generally less accessible in pure form than synthetic compounds. This is primarily due to their low abundance in natural sources and complex purification methods [22, 23]. Recent technological advances in analytical methods such as metabolomics, metabolic engineering and synthetic biology, as well as those in functional assays and phenotypic screens are opening new opportunities for natural compound-based drug discovery [2, 22, 24]. Increasing number of computational tools [25, 26], techniques [27, 28], and databases [29] are providing more accessible and powerful alternatives to explore the therapeutic potential of natural compounds.

An attractive approach to assess the bioactivity potential of a compound is comparing its chemical and structural similarity with that of the molecules with known activity [27, 30, 31]. Chemical structures encode complex atom and bond connectivity information which can be computationally exploited to predict their potential biological interactions. The chemical similarities between drug and natural compounds, especially dietary compounds, and their association with drug targets have been studied previously [12]. However, similarity assessments can vary considerably depending upon which structural fingerprint encoding is used [32, 33]. Indeed, the structural similarity between two molecules is a subjective concept [34] and no single similarity measure can likely capture the complex structure-activity relationships (SAR). Thus, owing to the vast structural diversity of natural compounds, it would be advantageous to include more extensive similarity measure encodings to more accurately establish structure-activity relationships and to predict bioactivities [27, 30, 35, 36].

Machine learning (ML) is being increasingly used to tackle complex structure-activity relations that are otherwise difficult to deconvolute [25, 37-39]. ML has been effectively used to predict, among others, molecular targets [40, 41], bioactivities [42], shared molecular interactions [43, 44], toxicity [45, 46], and drug-likeness of molecules [47]. Although ML models greatly facilitate predictions, they often lack interpretability which translates to the acceptance of their predictions in pharmaceutical or clinical settings. Thus, experimental validation assessing model prediction is crucial to build trust in a method [26].

Chemical proteomics (high-throughput methods for exploring drug-target-phenotype relationships) exists as a key and powerful method for target identification and elucidating mode of action of natural compounds [48, 49]. We propose a powerful approach where we combine computational target prediction of natural compounds with an in-vitro validation of the binding partner.

In this study, we identify potential of natural compounds (especially ingested dietary molecules) to bind human proteins that are known drug targets. We trained a binary machine learning classifier using chemical similarity scores from multiple fingerprints and physicochemical properties of paired small-molecule drugs with their known protein targets. The resulting model is then used to predict the molecular targets of hundreds of natural compounds through assessing their similarity with the drugs. We validate the model by demonstrating a predicted link’s Cox-1 binding activity by 5-methoxysalicylic acid (found in tea and herbs).

## Results

### Dataset of drugs with known targets

We utilized mappings between 1,410 FDA approved drugs (**Table** S1a) and their known, curated, targets (**Table** S1b) as our gold-standard dataset. The drugs were categorized according to their ATC (Anatomical Therapeutic Chemical) class, and into 16 chemical Superclasses (a hierarchy in chemical taxonomy with general structural identifiers such as organic acids and derivatives, organometallic compounds) [50] based on their chemical structures (Methods). Many of the drugs target the nervous system (264), followed by cardiovascular (180), anti-infectives (148), multiple ATC (131) and anti-neoplastic (127) (**Fig** S1a). Among the 16 structural classes, benzenoids and organoheterocyclics constitute the major super-classes of drugs (840) encompassing all therapeutic classes except the nutraceuticals (**Fig** S1a).

For the 1,410 drugs used, there were 1,262 known curated targets [51]. The number of drug targets ranged from 1 to 86 (i.e., some drugs have up to 86 known targets) (**Table** S1b), highlighting the fact that some drugs are well studied in terms of their target space. The most frequent targets were the different units and subunits of GABA receptors and GPCRs (adrenergic, muscarinic, histamine and dopamine receptors). The abundance of GABA receptors is consistent with the fact that many drugs (264) are targeting the nervous system.

### Predictor variables for representing a chemical pair

To deconvolute the complex structure-activity relation between drugs and their targets, we created a dataset for supervised binary classification. The underlying hypothesis is that a pair of compounds sharing at least one common protein target will be close in a high-dimensional structural space. A ML model for the latter is built using a binary classifier (one or more shared targets, or no shared target) with various structural similarity metrics as predictor variables. Such model can then be applied to predict the targets of natural compounds when compared to the drugs using the same predictors.

The first step towards building the classifier was to identify effective predictor variables. For this, similarity scores between all drug pairs were calculated using seven molecular fingerprints, viz., Morgan, Featmorgan, AtomPair, RDKit, Torsion, Layered and MACCS. The fingerprint similarity was scored using Tanimoto Score, which is measured on a scale of ‘0-1’; higher the score, more similar are the molecules. Consistent with each molecular fingerprint assessing different features of the compound, the tanimoto score distribution for drug pairs differed across all the seven fingerprints (**Fig** 1B). For most of the drug pairs, the Morgan, Featmorgan and Torsion fingerprints consistently yielded a lower score (tanimoto score_median_ = 0.11, 0.13, 0.08 respectively) as compared to AtomPair, RDKit, Layered, and MACCS (tanimoto score_median_ = 0.23, 0.36, 0.45, 0.32 respectively). The drug pairs also showed broader tanimoto score distribution in AtomPair, RDKit, Layered and MACCS. A rank comparison of drug-pairs (**Fig** S2) showed a low concordance amongst the different fingerprints supporting that each fingerprint captures different aspects of the structural similarity. This variance, together with the previous observations that none of the fingerprints alone is universally suited [32, 52], we decided to utilize the similarity scores from all the seven fingerprints in training the ML classifier.

**Figure 1.**
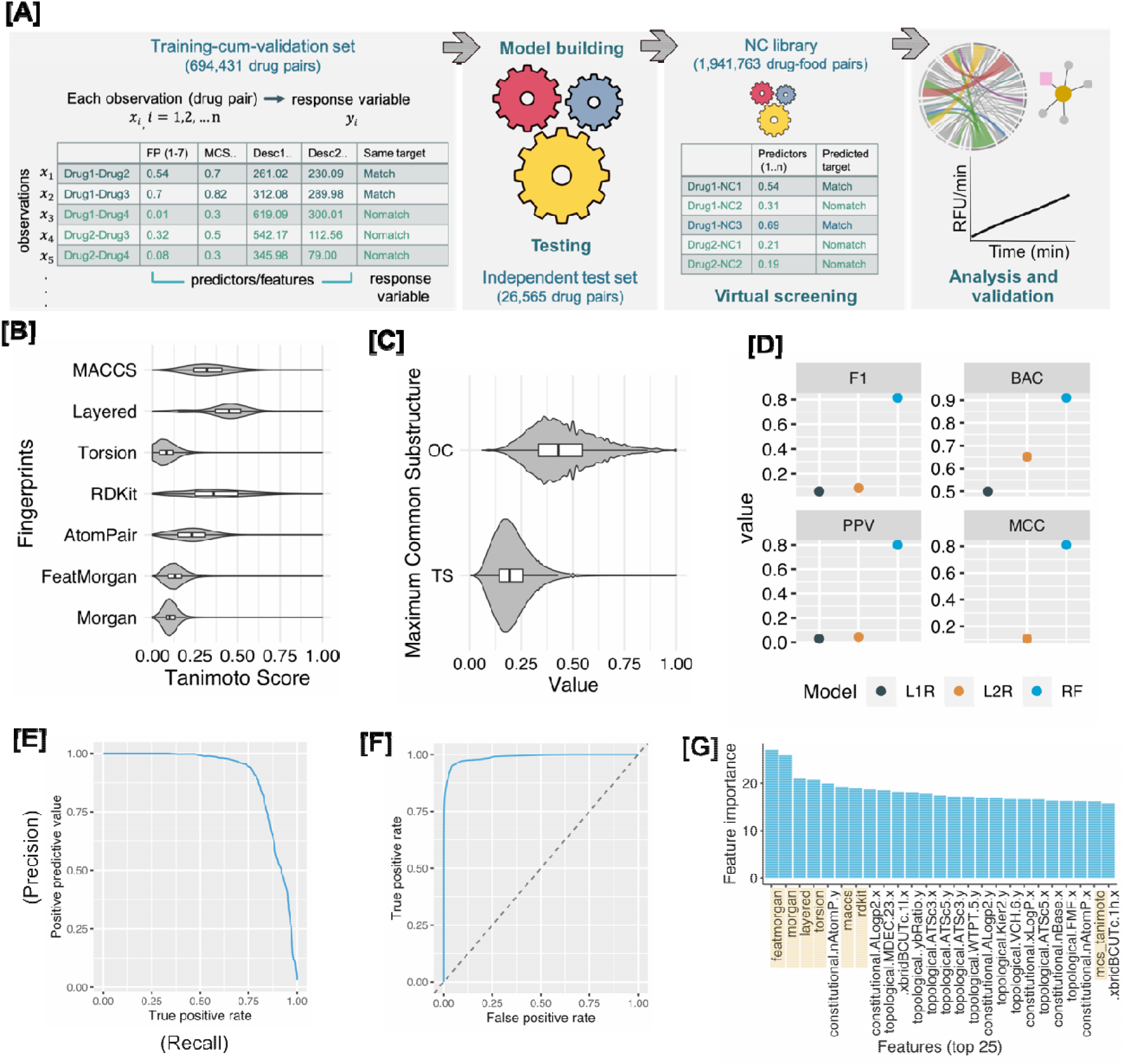
(A) **Overview** of workflow deployed. A training-cum-validation set comprising of drug pairs was created using various predictor variables (fingerprints, MCS and physicochemical properties). The model was trained for response variable (Match or Nomatch) and tested on an independent test set for performance evaluation. The natural compound library paired with drugs was virtually screened to obtain hit pairs, followed by analysis and in-vitro validation. (B-D) - **Similarity metrics (ML dataset)**. (B) Molecular fingerprints - the 7 fingerprints generate a different similarity score for the pairs of drug molecules compared. The median value of each is represented in the box plot (in the center) and the spread shows the density of the drug pairs around that score. (C) MCS - there are two types of scores reported by the MCS algorithm, one is the Tanimoto score and the other is the Overlap coefficient (OC). The violin plots were smoothed for density by an adjustment factor of 3. (D-F) - **Performance on the test set**. (D) performance of the three models namely regularized logistic regression (L1R and L2R) and random forest (RF) on independent test set. Performance was evaluated using balanced measures: F1 score, balanced accuracy (BAC), positive predictive value (PPV) and matthews correlation coefficient (MCC). RF clearly had higher performance as compared to the logistic regression models under all metrics. The hypertuned RF model was selected for further evaluation using (E) precision-recall and (F) ROC curve – the model achieved a high AUC of 0.98 on the independent test set. (G) High ranking features of RF model - top 25 features are displayed, showing most of the distance-based features provided maximum information gain with ‘Featmorgan’ performing best.

In addition to the fingerprint metrics, maximum common substructure (MCS, which is based on overlap between the two molecules represented as chemical graphs) [53] and the physicochemical descriptors (numerical properties) were included as additional predictor variables based on their previously noted utility [25, 54, 55]. The MCS calculation reports several statistics, amongst which the MCS size (median = 8), tanimoto score (median = 0.19) and OC (overlapping coefficient) (median = 0.43) score are important measures to assess similarity. The tanimoto score and OC score distribution is shown in **Fig** 1C. OC, measured on a scale of 0-1, accounts for size difference amongst molecules and is a useful indicator when there is a significant size difference between molecules being compared. The MCS measures are more intuitive to interpret as the substructure graph shared between the two molecules can be visualized (as in Fig 3A) and can be mapped back to the underlying molecules to extract which are the common and unique features, while this is not possible with the fingerprints and the molecular descriptors.

For molecular descriptors, we used 5 different categories (constitutional, topological, geometrical, electronic and hybrid) to capture individual physicochemical properties of the drugs. **Table** S1d, reports the number of descriptors used in each category. Majority of the physical and chemical information comes from the constitutional and topological descriptors. In total 225 molecular descriptors (for example, molecular weight, logP, aromatic bonds, and ring blocks) were calculated for each molecule.

### Data Processing

All pairwise distance-based measures (tanimoto score of Morgan, Featmorgan, AtomPair, RDKit, Torsion, Layered and MACCS), MCS features (MCS size, MCS tanimoto score, MCS overlap coefficient), and the molecular descriptors (constitutional, topological, geometrical, electronic and hybrid) were merged to create a matrix of 460 predictors and a binary response variable (‘Match’ or ‘Nomatch’).

#### Training and test set

Prior to training the classifiers, the data was split into a training-cum-validation set (80%) and a hold-out test set (20%). The splitting might result in overlapping predictors (i.e., the physicochemical descriptors) in train and test set as the drugs are paired. Thus, in order to avoid overlapping drug predictors, we adopted a systematic stratification procedure to split the paired drugs to ensure that the train and test sets were independent. This would essentially mean that all the drug pairs present in the test set were exclusive to it and were not seen by the classifier during learning from the training set. This procedure ensured that the classifier did not depend on availability of exact drug pairs in the test to make predictions and a high accuracy would indicate generalization to novel drug pairs. In order to achieve this, we performed a random selection of drugs using their Superclass information (as some drug classes are over/under-represented, **Fig** S3) to have a balanced representation. We randomly selected 20% drugs from the highly represented drug Superclasses (i.e., at least > 100 drugs, Fig S3) for our exclusive test set. This stratification resulted in 1,179 drugs (694,431 pairs) in the train set and 231 drugs (26,565 pairs) in the test set.

#### Data pre-processing

The data was preprocessed to remove non-informative features by removing constant variables such as with all 0’s or all 1’s (*n=82*), and to filter out observations with missing values (*n=23,370*) for any predictor variable. This resulted in a dataset with 378 predictors and 671,061 observations. The classification models were then trained to classify the binary response variable for target ‘Match’ (*n=19,542*) and ‘Nomatch’ (*n=651,519*).

### Machine Learning

We employed two learning algorithms commonly used for binary classification tasks for large datasets, namely, logistic regression (LR) [56] and random forests (RF) [57]. This selection was done to cover models that can capture linear (LR) and nonlinear (RF) relationships as both provide interpretability in terms of feature weights as well as probabilistic predictions.

#### Logistic regression

Logistic regression models apply logistic function to weighted linear combinations of their input predictors to obtain model predictions. We used regularized logistic regression (L1R and L2R) so that the overall error (cost function) during training is minimized to optimize performance while controlling the complexity of the models leading to better generalizability [58]. The models were trained for optimized learning by setting the cost parameter C*= 9*.*730296e-11*. This value was determined using a heuristics on a balanced subset of data [59]. To account for the highly imbalanced nature of the training set we applied class weights (Match – 0.97, Nomatch – 0.03). Both regularization types, L1R and L2R, were used to train the data and their performance on the hold-out test set was evaluated (**Fig** 1D). As can be seen from **Fig** 1D, both the linear models failed to adequately predict the response variable suggesting higher order of complexity that cannot be segregated by a linear model.

#### Random Forest

A random forest consists of many individual decision trees, each of which is trained on a subsampled version of the original dataset. The predictions of individual trees are averaged to provide a final prediction for the full forest. Classically, RFs are known to have strong performance on various computational chemistry tasks and have been state-of-the-art in various cheminformatics settings till date such as to predict chemical binding similarities [35], learning drug functions from their chemical structures [60], in-vitro toxicity prediction [61], and drug-target interactions [62].

Since the default parameters might not be optimal for any learner, we used hyperparameter tuning for building an optimized RF model. Hyperparameters (ntree (number of trees), nodesize (number of observations at terminal nodes), mtry (number of variables to split at each node) and classwt (class weight) were randomly searched over 10-iterations in a 5-fold cross-validation setting (**Fig** S4). It generated 10 combinations of tuned parameters from which the final parameters were selected by the learner using the performance metric MCC [63]. Even with other commonly adopted metrics such as F1 (**Fig** 1D) or accuracy (**Fig** S4), the set of best performing hyperparameter combination was consistent. The hyperparameter tuned results (**Fig** S4) showed all 10 iterations performed comparably indicating robustness of the RF on our dataset. The best performing hyperparameters combination – *ntree=241, nodesize=27, mtry=23, classwt=2,515* (selected by best value of MCC) – was used to train the final model. The model was tested using the independent test set and performance metrics are depicted in **Fig** 1D-F. Data skewness can significantly impact performance metrics [64]. We therefore used a combination of metrics to evaluate the model’s performance on the test set. We focused on balanced threshold metrics (**Fig** 1D) (matthews correlation metric (MCC) – 0.81, F1 – 0.82, Balanced accuracy (BAC) – 0.911) in combination with rank metrics (**Fig** 1E and 1F) (ROC and precision-recall curves) to evaluate our model’s performance (metrics definitions explained in **Table** S3). Overall, the RF model performed robustly, achieving an AUC of 0.988, precision of 0.80 and recall of 0.83.

#### Extracting feature importance

During training, RF models were configured to generate feature/variable importance measures whereby the ‘MeanDecreaseGini’ (MDG) is measured, which is based on the Gini impurity index used for the calculation of the decision tree nodes. MDG is the total decrease in node impurities from splitting on the variable, averaged over all trees [65]. We extracted these feature importance values from the final trained model to explore which predictors contained highly predictive information. The feature importance from high to low (only top 25) are shown in **Fig** 1G. The ‘Featmorgan’ was the best predictor variable with the highest importance value in our trained model closely followed by other distance-based fingerprints.

### Predicting natural compounds and drugs similarities

#### Source of natural compounds and creation of drug-food pairs

A catalogue of 11,788 natural compounds was obtained from FooDB (www.foodb.ca) (**Table** S1c). These correspond to 261 unique food sources and are categorized into 15 main food types such as vegetables, fruits, herbs and spices, and milk products (**Fig** S1b). For the simplicity in representation in **Fig** S1b, the frequency accounts for only one source per compound; however, a particular compound can be present in multiple food sources. The food compounds were structurally classified into 21 classes (see methods) (**Fig** S1b). Highly represented were lipids and lipid-like molecules (4803), phenylpropanoids and polyketides (2476), organoheterocyclics (1381) and organic oxygen compounds (1120). All these natural compounds were used to create an assessment library, where each chemical pair comprised of a drug and a natural compound (**Fig** 1A) (now referred to as drug-food pair). Pairwise similarities between each natural compound and all the drugs were computed using the same set of predictors (i.e., using same predictors molecular fingerprints, MCS and molecular descriptors) as was for training dataset.

#### Similarity predictions by RF model

The similarity of the 11k natural compounds paired to 1410 drugs was evaluated using the trained RF model. As many drugs have originated from natural compounds, drug-food pairs with a very high similarity fingerprint score (i.e., tanimoto score > 0.9) were removed (n = 1,850 pairs) considering them to be the same compound (which also serves as a validation of the model). Overall, the number of drug-food pairs compared were 1,941,762. 574 natural compounds passed the threshold of 0.5 probability of being considered as a match to at least one drug. The number of hits at different probability thresholds are shown in Fig 2A. Probability threshold of 0.6 (194 pairs) was used in further analysis.

**Figure 2.**
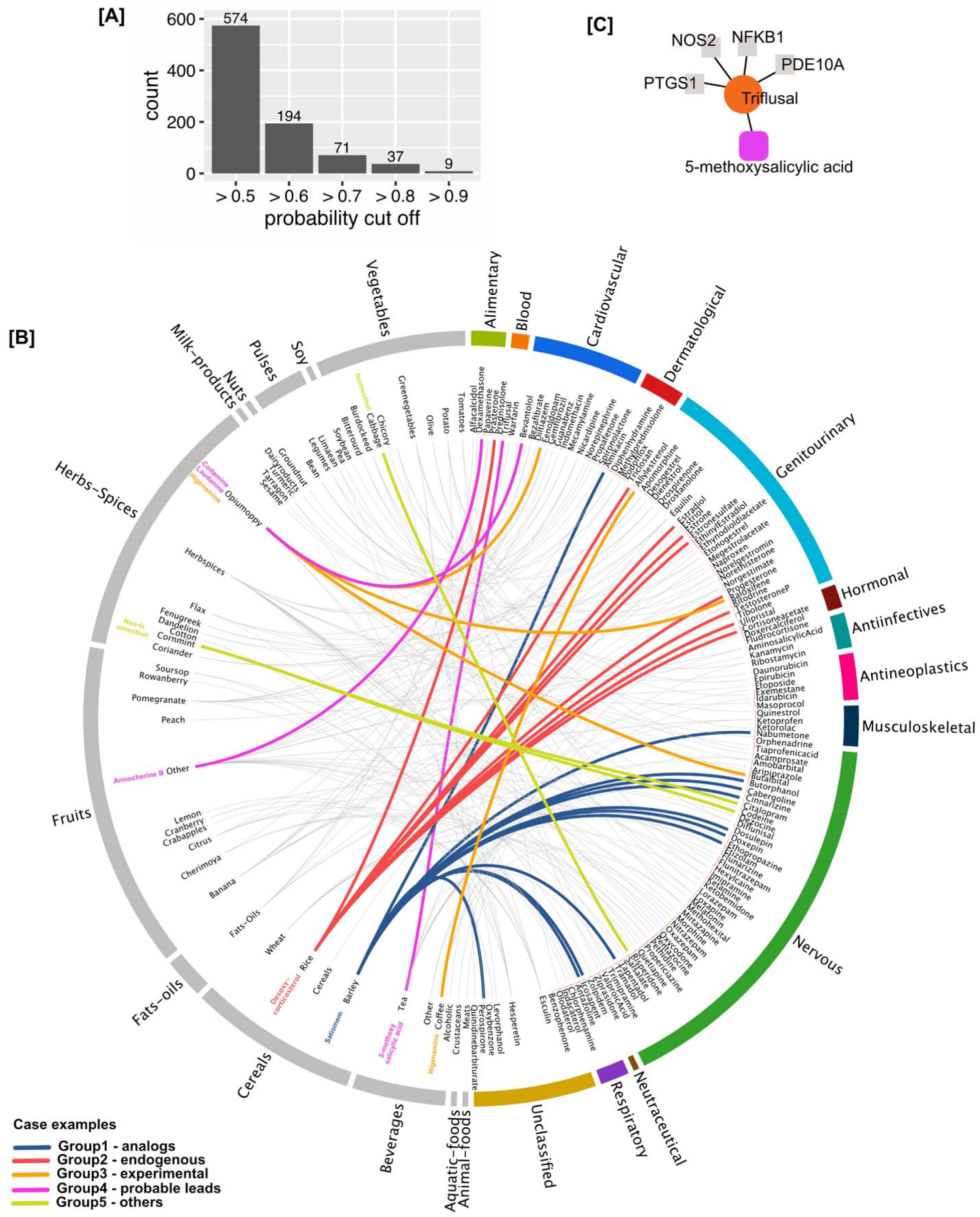
Drug-food similarity. (A) Number of hits at different probability thresholds. We chose probability > 0.6 to report the results. (B) 194 drug-food pairs predicted as ‘match’ at the probability threshold of >0.6. The drugs are arranged according to their therapeutic class and food compounds according to their food source. The highlighted colored links represent the case examples in the five author defined groups (details in the text). (C) Group4-probable lead example taken up for experimental validation. The food compound 5-methoxysalicylic acid was a hit with the drug triflusal which has 4 known targets. We validated the inhibitory activity of triflusal and 5-methoxysalicylic acid against the target PTGS1 (also known as Cox-1).

The 194 drug-food similarities (**Fig** 2) comprise of 107 unique food compounds and 136 unique drugs (full annotated list in **Table** S6). Note that a drug can share similarity with more than one food compound and vice-versa. Also, a food compound can be present in multiple food sources, for exhaustive listing of known sources, we recommend querying the FooDB using respective compound Ids or names.

We performed manual curation of these 194 drug-food pairs and categorized the food compounds into five custom defined groups based on the meta information available in the public domain (**Table** S6, Fig 2B). Group 1, Analogs: food compounds closely resembling another known drug available in the market. Group 2, Endogenous: compounds reported as a metabolite in humans. Group 3, Experimental: food compounds currently under investigational or clinical trial as a therapeutic. Group 4, Probable lead: compounds with potentially novel bioactivity. Group 5, Others: compounds currently used in industrial application or used as additive or flavor enhancer. Each group is discussed below with case examples.

##### Group 1-analogs (51 pairs, 17 natural compounds)

food compounds in this group were found to be analogous to other known drugs. This is consistent with the natural compound origin of many drugs, and thus this group served as a control illustrating that the model captured meaningful relations. Yet, presence of some of these compounds in food is intriguing. A compound referred to as ‘Satiomem’ in FooDB (reported in barley and onions) resembles the drug Carbinoxamine, an antihistamine. It shared similarity with other antihistamines (such as Chlorprothixene, Doxepin, Antazoline and Chlorphenamine, **Fig** 2B, highlighted in blue) belonging to the nervous ATC category which are used as antipsychotics. These drugs share ‘Histamine H1 receptor’ as a target but there was no evidence found for Satiomem/Carbinoxamine having antipsychotic activity so it could potentially serve as a good candidate for further testing as an antipsychotic or resulting in drug interactions when used in combination with these drugs.

##### Group 2-endogenous (54 pairs, 24 natural compounds)

compounds that are endogenous to human tissues but also reportedly present in various food sources. For example, ‘desoxycorticosterol’ a.k.a. 21-Hydroxyprogesterone was reported to be present in rice and is endogenously present in amniotic fluid and blood throughout human tissues. It’s predicted to be similar to other Hormonal and Genitourinary drugs (**Fig** 2B, highlighted in red). ‘Estriol’, an estrogen produced by the human body, is reported to be present in pomegranate and beans.

##### Group 3-experimental (9 pairs, 6 natural compounds)

these food compounds are already under experimental investigation category (i.e., under approval to be used as drugs, reported in DrugBank accessed January 2018). ‘Higenamine’ is reported to be present in opium and coffee. This compound is in clinical trial (DrugBank id: DB12779) and has been patented for various therapeutic applications (**Fig** 2B, highlighted in orange). This group serve as a proof of principle that we could recall natural compounds with similar activity as currently used human-targeted drugs, which are being actively investigated pre-clinically.

##### Group 4-probable lead (72 pairs, 55 natural compounds)

to our knowledge, the compounds in this group have little or no hitherto reported evidence of their physiological or biological activity. The drug Papaverine is an alkaloid which is a vasodilator. ‘Annocherine B’ reportedly present in many fruits showed high similarity with Papaverine, however no evidence or reports of this compound about its action or use was found. ‘Bevantolol’ is a cardiovascular drug which shared similarity with two novel compounds ‘Codamine’ and ‘Laudanine’ (both reported in opium).

##### Group 5-others (8 pairs, 5 natural compounds)

food compounds found in this group are reported to be used as food additives such as flavor enhancers or have other industrial applications such as emulsifiers. ‘Neoisomenthol’ and ‘Isomenthol’ are used as a flavoring agent are similar with nervous category drugs ‘Codeine’, ‘Dezocine’, and ‘Tapentadol’. Thus, these five groups highlighted interesting similarity relationships existing between drug and food compounds and their wider therapeutic potential.

### Experimental validation of Cyclooxygenase-1 (Cox-1) inhibition by 5-methoxysalicylic acid

With the growing number of new and improved learners [66, 67] there’s always a possibility to benchmark and compare the performances with different learners. There are multiple scenarios under which one learner can be chosen over another for a given problem. For instance, the performance of nonlinear RF model over the linear learner logistic regression models has been benchmarked previously on a large number of experimental datasets [68]. While this complements our observation, our major goal was to present a proof-of-concept of the learner’s predictions in real and not merely restrict to their performance comparisons. As has been proposed, complementary experiments that can validate any model’s predictions can help build trust in the method or it’s outputs [26].

An interesting case for experimental follow-up in our 194 high-confidence similarity pairs is that of triflusal and 5-methoxysalicylic acid (**Fig** 2C). Triflusal is an antithrombotic anticoagulant and is considered very important for the secondary prevention of ischemic stroke. Triflusal has an antagonist effect on prostaglandin G/H synthase 1 (PTGS1) (also called Cox-1) in platelets [69]. Consistent with our prediction, 5-methoxysalicylic acid, which is found in tea, herbs and spices, has been shown to have antiplatelet activity in rats [70]. Yet, the molecular target of 5-methoxysalicylic acid is not known. We therefore chose to assess Cox-1 binding activity of 5-methoxysalicylic acid. To attest the utility of ML approach in comparison to using a single similarity measure (such as a selected fingerprint), we also tested a ‘negative control’ molecule, 4-isopropylbenzoic acid, which is found in cumin, herbs and spices. This molecule had the highest similarity with triflusal based on Featmorgan, which was the most important predictor (**Fig** 1G). While 4-isopropylbenzoic acid was predicted as ‘Nomatch’ by the RF model, our test compound (5-methoxysalicylic acid) was predicted as a ‘Match’ but had a lower Tanimoto Score for Featmorgan (**Fig** S5A). Structural representation of all the tested compounds and their shared MCS (maximum common substructure) is depicted in **Fig** 3A.

**Figure 3.**
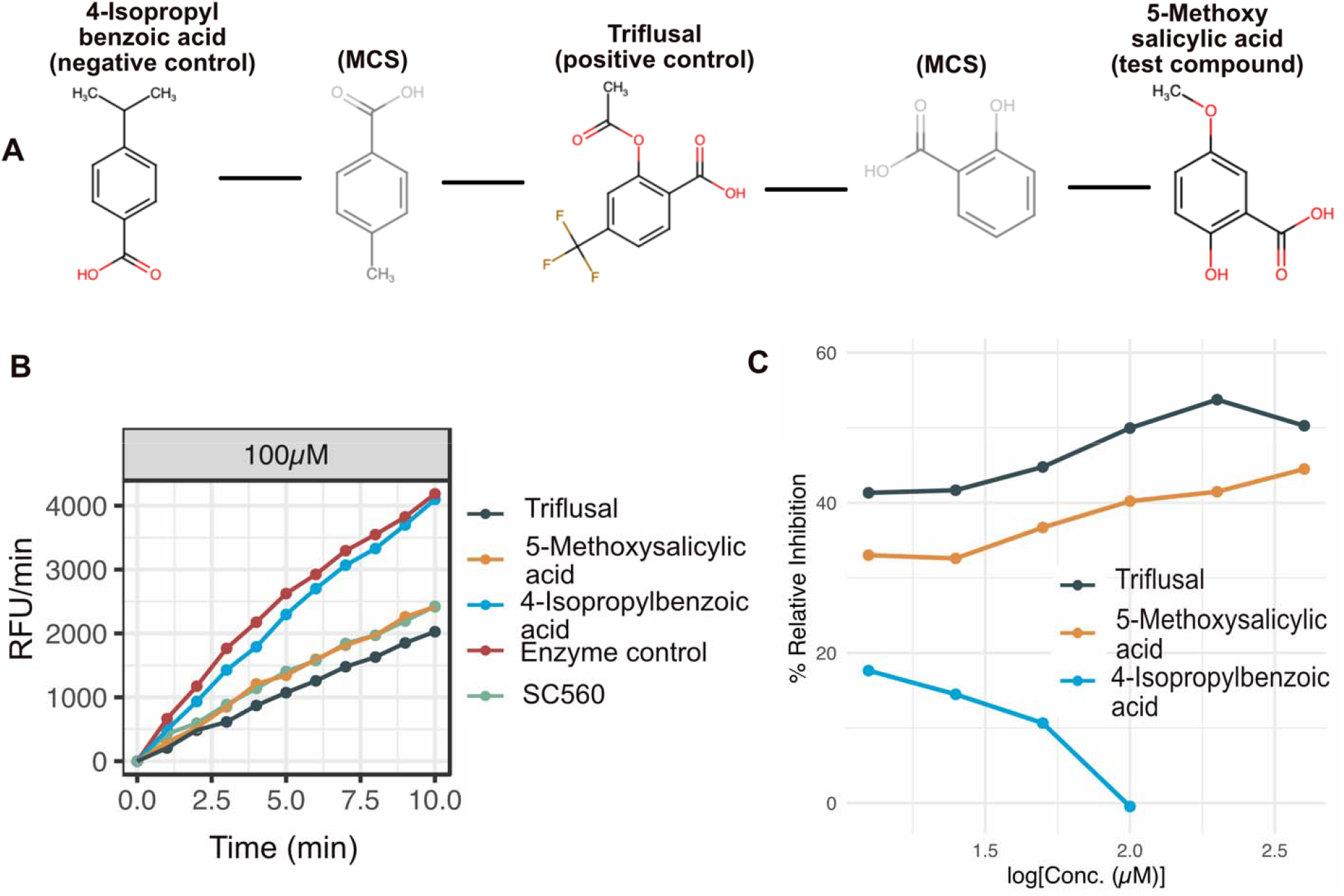
Cox-1 inhibitor assay. (A) Chemical structures of all the tested compounds. MCS structures are also depicted which helped to intuitively assess the structural similarity between the tested compounds (B) An example relative fluorescent units (RFU) plot of the tested compounds at 100μM (other tested conc.: 12.5μM to 400μM serial dilutions). SC560 is a positive control provided by the assay kit supplier (Methods). (C) Relative inhibition of the positive control (drug triflusal), test compound (5-methoxy salicylic acid) and negative control (4-isopropyl benzoic acid) at different tested concentrations. 5-methoxy salicylic acid showed similar inhibition of Cox-1 as the drug triflusal whereas no such inhibition was observed for 4-isopropyl benzoic acid. 4-isopropyl benzoic acid showed strong color change (bright pink) reaction beyond 100μM and thus was found unsuitable for being tested at higher concentration with this assay.

We tested the Cox-1 inhibitory activity of the three compounds, triflusal (positive control), 5-methoxysalicylic acid (test compound), and 4-isopropylbenzoic acid (negative control) using an enzymatic assay based on fluorometric detection of prostaglandin G2, which is an intermediate product generated by the Cox enzyme (Fig S5B).

Triflusal and 5-methoxysalicylic acid exhibited highly similar inhibition profiles over different concentrations (Fig. 3B, Table S4). The IC50 (is a measure of potency of a substance in inhibiting a biological or biochemical function) of triflusal was estimated to be ∼100 μM, while 5-methoxysalicylic acid showed ∼40% inhibition at 100μM. Maximum inhibition achieved with SC560 – a positive control included in the assay kit – was 42%, comparable to that of 5-methoxysalicylic acid. In stark contrast, 4-isopropylbenzoic acid did not show any inhibition at 100 μM. At higher concentrations, it caused a color formation (bright pink) and hence was not tested beyond 100 μM. It only showed a small (10-17%) inhibitory effect at a lower concentration, possibly be due to non-specific binding. Taken together, the ML predicted drug-natural compound pair experimentally showed the same target binding with quantitatively similar activity, supporting the underlying model.

### ML vs single fingerprint

In order to compare the performance of using a single fingerprint alone (‘featmorgan’ as it was the best predictor in our analysis and is also popularly used) versus our RF model trained in this study, we used different thresholds of fingerprint similarity to capture the hits.

Featmorgan based hits grew significantly in number as the tanimoto score threshold was lowered (> 0.7 – 21 compounds, > 0.6 tanimoto score – 112 compounds, > 0.5 – 451 compounds). This behavior is not helpful in a drug discovery setting where computational shortlisting is meant to reduce the number of hits that can be thereafter tested experimentally. The validation pair triflusal-5-methoxysalicylic acid was predicted as a match with high probability by ML model (0.72) but had a much lower tanimoto score for Featmorgan (0.36). This pair could have been easily missed (would be a false negative) if Featmorgan was used alone to rank molecule similarity and thereby estimate their activity profile.

## Discussion and Conclusion

This study as addressed three goals: identification of potential molecular targets of ingested natural compounds and exploring their therapeutic potential; evaluating the utility of a comprehensive ML based approach to deconvolute the complex SAR between molecules as opposed to restricting to a single similarity measure; and, lastly, complementing in-silico model predictions with experimental validation to build trust in model’s predictions.

Systematically integrating computational chemistry approaches can help deconvolute the intricate structure-activity relationship between small molecules and their biological targets. The data fusion approach – i.e., integrating multiple similarity metrics based on fingerprints, maximum common substructures, and physicochemical descriptors – used in this study proved effective in identifying natural compounds functionally similar to known therapeutic drugs. The RF model achieved a high performance with balanced accuracy of 0.911 and MCC of 0.81. Analysis of 194 drug-food pairs predicted from the model helped to capture drug analogs, host endogenous metabolites, some investigational drugs, as well as novel molecular leads present in various food sources which are deemed to share the same target as the drugs.

Data fusion approaches can aid in reducing the amounts of artefacts [30]. In line with this, various methods are being proposed to accelerate the performance of in-silico similarity searches such as inclusion of bioactivity profiles [27, 71, 72], multiple fingerprint algorithms [73], and similarity ensemble approach [74]. Our approach serves as another promising addition to these strategies. A recent study has also demonstrated similar strategy to identify molecules using pairwise similarity concept and target engagement [35] which supports our strategy and showcases the utility and success rate of such methods in early-stage molecule discovery.

In the context of natural compounds search space, two closely related issues remain open before the proposed approach could be more broadly applied: the estimation of true-positive rate, and the availability of curated molecular target information for an additional, structurally more diverse, set of small molecules. The former could be addressed through high-throughput testing of bioactivity profile of natural compounds against a set of protein targets using cell-based or enzymatic assays [75, 76]. The generated data could then be used to address the latter issue. Further, new structured and curated datasets such as those recently reported by Duran et al [27] would be valuable to this end.

The ML approach used here was notable in capturing complex high-dimensional similarity that would not be accessible based on any structural similarity metric used in isolation. Indeed, we could show that a molecule that is highly similar based on a single fingerprint’s tanimoto score did not show any appreciable activity. In contrast, the natural compound identified using RF model trained on multiple features showed the predicted enzyme inhibitory activity. In further support, the model could also identify several drug-food compound relations including compounds that are currently under investigation or have been ascribed with related bioactivity in the literature. Taken together, this study has implications for efficient exploration of drug-like properties of natural compounds.

## Materials and methods

### Data Source and processing

All the FDA approved drugs which had target information (**Table** S1a and S1b) associated with them were taken from DrugBank [51] (accessed January 2018). The natural compound library used for virtual screening was obtained from FooDB (www.foodb.ca; freely available and accessed June 2017, (∼11k compounds). It was curated and formatted in order to be smoothly integrated into our analysis. The full list of compounds with their annotations is provided in **Table** S1c. It included compounds from both raw and processed foods. We used drug classification codes from ATC (https://www.whocc.no/) to therapeutically classify all the drugs and ClassyFire [50] to structurally classify the drugs and the natural compounds.

### Computing predictor variables

In order to predict the molecular targets for the natural compounds based on their pairwise structural similarities with the drugs, we needed to create a ML dataset that contains molecular features (i.e., predictors) computed from the chemical structures of the drugs and gives out a binary response for the target. This model can turn then be applied to natural compounds and drug pairs to predict the potential targets for natural compounds. For this, a number of pairwise distance measures and molecule specific physicochemical descriptors were generated for drugs and all the natural compounds. These included distance-based fingerprint similarities, maximum common substructure similarities and physicochemical descriptors. These molecular features formed the basis of our ML dataset (explained in later section) created from drugs and their known respective targets.

Fingerprint calculation, Tanimoto Score estimation and Molecular descriptor generation

The INCHIs from drugs and natural compound library were used to generate 2D structural information in **S**tructural **D**ata **F**ormat (SDF). These SDF files were used to calculate 7 different molecular fingerprints (Morgan, FeatMorgan, AtomPair, RDKit, Torsion, Layered and MACCS) to gather theoretical 2D structural information from the molecules. The pairwise structural similarity from the fingerprints was scored using the widely used Tanimoto similarity metric (computed as *TS*_*AB*_ = (*A* ⋂ *B*)/(*A* ⋃ *B*)). The entire workflow was designed using the KNIME [77] analytics platform utilizing RDKit [78] plugin (for fingerprints) with default parameters. We also computed maximum common substructure (MCS) shared between each chemical pair. The MCS is a graph-based similarity search wherein the largest substructure shared between query and target is identified and gives out various parameters such as number of MCS generated, size of each molecule, size of MCS, tanimoto score, overlapping coefficient (computed as *OC*_*AB*_ = (*A* ⋂ *B*)/*min* (*A, B*)). It was computed using the ‘ChemmineR’ [79] and ‘fmcsR’ [53] packages available for R. Although MCS calculation is computationally intensive and time-consuming, but they are more sensitive, accurate and intuitive, thus we implemented this computation in batch-mode on high-performance computing cluster for faster processing. In addition to the distance-based features, five different types of molecular descriptors (constitutional, topological, geometrical, electronic and hybrid) were computed for all compounds using R package: RCDK [80].

### Data preprocessing and Machine Learning

All predictors (computed above) for the paired drug data were combined into a single matrix with each observation associated with a response variable (‘Match’ or ‘Nomatch’). This data was then split into an 80% train-cum-validation set and a 20% test set by performing random selection of drugs from the highly represented Superclasses such that both sets were mutually exclusive and had a balanced distribution of the drug classes. Following this the training set was pre-processed to remove constant predictors and missing value observations to avoid training on noise.

We explored both linear and nonlinear learners to build our binary classification models. The linear learners used were the two types of regularized logistic regression: called as L1R and L2R (‘LiblineaR’ package in R) and the nonlinear learner was random forest (‘mlr’ package in R). The binary classifiers were trained for two classes which are referred to as ‘Match’ and ‘Nomatch’ indicating whether they share a target or not. In order not to increase the system complexity in terms of protein target similarity (as drugs can share multiple targets), each drug pair which shared even at-least one target were considered as ‘Match’ and the rest were considered as ‘Nomatch’. Modern machine learning algorithms require tuning various parameters in order to achieve their best performance [81]. Thus, the classifiers used were optimized accordingly by tuning their parameters.

L1R and L2R logistic regression are extremely fast learners and requires input data to be centered and scaled in a *nxp* numerical matrix form and a response variable (*1xn*) containing class labels which can be a vector or factor. The two types of regularization used were L1 (type 6) and L2 (type 0) of LiblineaR package which can give out probability estimates of prediction. As our paired data was highly imbalanced, we used class weights where the positive class received a higher weight ratio (Match - 0.97) then the negative class (Nomatch - 0.03). The weights were derived from ratio of positive/negative classes. To handle misclassification errors, costs were determined by using ‘heuristicC’ function on balanced sub-sample of the dataset [59].

We also defined a search space to tune the hyperparameters for random forest (RF) to boost its performance. Four different and broadly used hyperparameters were tuned: the number of trees (*ntree*), the number of observations at terminal nodes (*nodesize*), number of variables to split at each node (*mtry*) and class weights (*weight*). We limited the *ntree* to 300 as setting higher *ntree* values didn’t resulted in any significant difference in performance rather only increased the computational overhead (i.e., training time and memory usage). The default value for the parameter *nodesize* is 1, but with low values of tree depth, the tree can fail to recognize useful signals from the data. We searched for *nodesize* value in the range 20-50. Lower *nodesize* can result in lower detection signals of the true positives and a high false-negative rate. The default 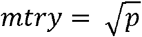 where p is the number of features in the input data. In our ML dataset, the default *mtry* was 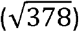 19 features. We searched *mtry* in the range of 15-30. Lastly, as our training data set had high class imbalance (Match-Nomatch ratio of 0.03), we also tuned the class weight parameter starting with a minimum weight of 300 for the positive class (‘Match’) and searching up to 10 times i.e., 3000. The hyperparameters were additionally optimized for 10 random iterations with 5-fold cross-validation each using stratification. Owing to the size of the data and to speed up the iterations and parameter search the tuning was performed in a cluster environment with parallel backend.

We used the standard performance measures (mean test values for MCC (matthews correlation coefficient), Balanced accuracy (BAC), Kappa, MMCE (mean classification error), ACC (accuracy), TPR (true positive rate/Recall/Sensitivity), FPR (false positive rate), TNR (true negative rate), (false negative rate), PPV (Precision/Positive predictive value)) to evaluate the learner’s performance in each iteration and model assessment.

### Compound preparation and Assay protocol

Briefly, all tested compounds were dissolved in their respective solvents. Triflusal and 5-methoxysalicylic acid were dissolved in DMSO and 4-isoproplybenzoic acid was dissolved in ethanol. Compounds supplied with the assay kit were prepared as per the manufacturer’s protocol and the assay was also performed according to the instructions present in the kit from Abcam (CAT#ab204698). The kit included the Cox-1 enzyme (source: ovine) and had a positive control Cox-1 inhibitor (SC560).

Literature evidence showed that triflusal binds to purified Cox-2 at 240-320 μM [82]. Thus, we assayed all compounds at different concentrations starting at 400μM and going down to 12.5μM with serial dilutions and in triplicates. Relative Fluorescence Units (RFU) were measured immediately after starting the reaction by using microplate reader (Tecan infinite M1000Pro) at Ex/Em = 535/587 nm in a kinetic mode for 40 minutes at 25° C. All fluorescence readings for triplicates under a given concentration were averaged and initial time point RFU reading was used to shift the measurements to start from 0 (**Fig** 4B). We took the first 10 time-points of RFU readings to assess the inhibitory effect of the tested compounds. Slopes for all samples (triflusal (positive control), 5-methoxysalicylic acid (test compound) and 4-isopropylbenzoic acid (negative control)), enzyme control, and kit supplied positive control (SC560) were calculated by fitting linear equations, respectively. Percent relative inhibition for samples was calculated as

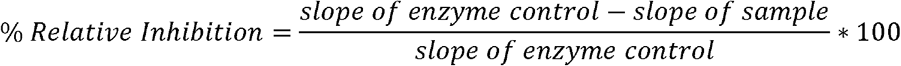

**Fig 4.**
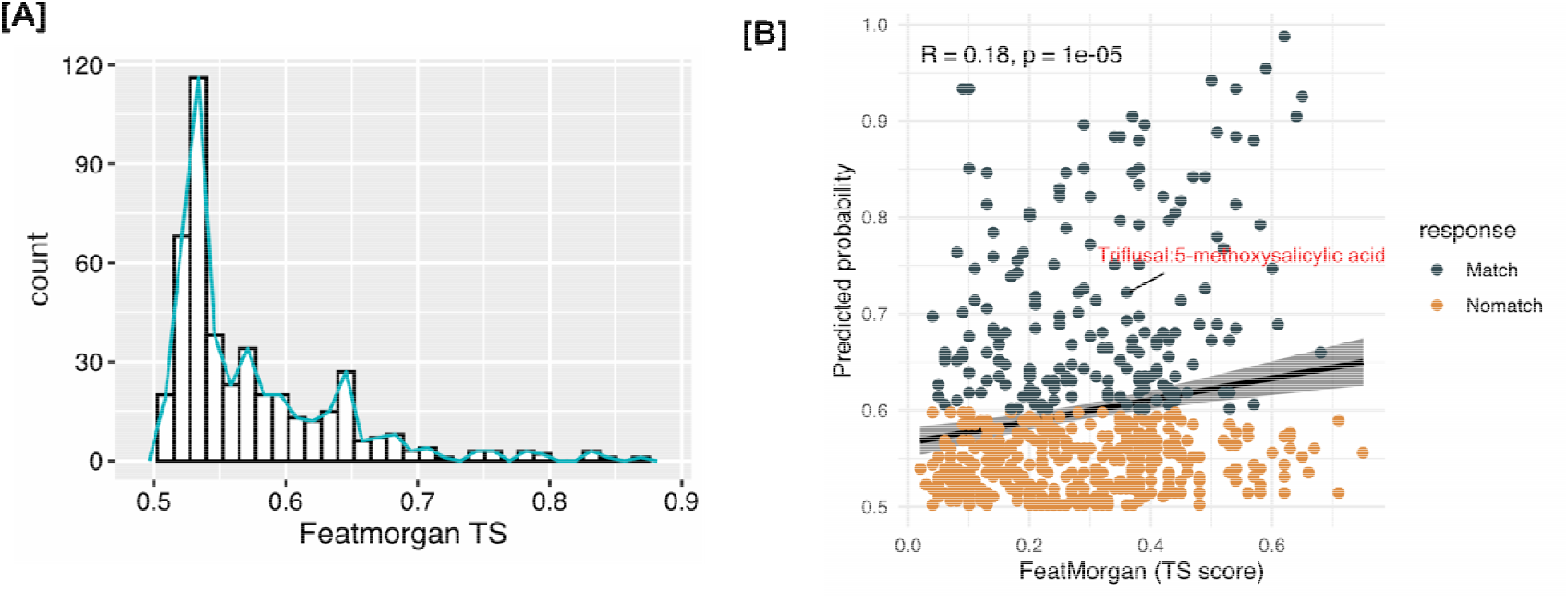
RF vs featmorgan. (A) number of hits retrieved by using tanimoto score of featmorgan as similarity measure, which grows significantly as threshold is reduced (lower threshold means less similarity). (B) correlation between RF probability predictions >0.5 (a pair is called match at probability > 0.6) with corresponding tanimoto score of featmorgan of drug-food pairs. Our hit pair triflusal and 5-methoxysalicylic acid (highlighted in red) was predicted a hit by RF model would be missed by featmorgan if used alone.

## Data and Code availability

The input data files can be accessed at (DOI: <u>10.17632/7ft539gwf3.1</u>) and codes for analysis and figure generation are available at https://github.com/periwal45/periwaletal2020.

## Acknowledgements

VP and NG were supported by the EMBL Interdisciplinary Postdoc (EI3POD) program under Marie Sklodowska-Curie actions COFUND (grant number 664726). SB was supported by the Add-on fellowship for Interdisciplinary Science of the Joachim Hertz Foundation.

