## Supplementary material for "Bioactivity assessment of natural compounds using machine learning models based on drug target similarity": Suppl. Fig. S1-S5 & Suppl. Tab. S3-S4

**Fig S1**: Structural classification of drugs (S1a) and foods (S1b).

**S1a**

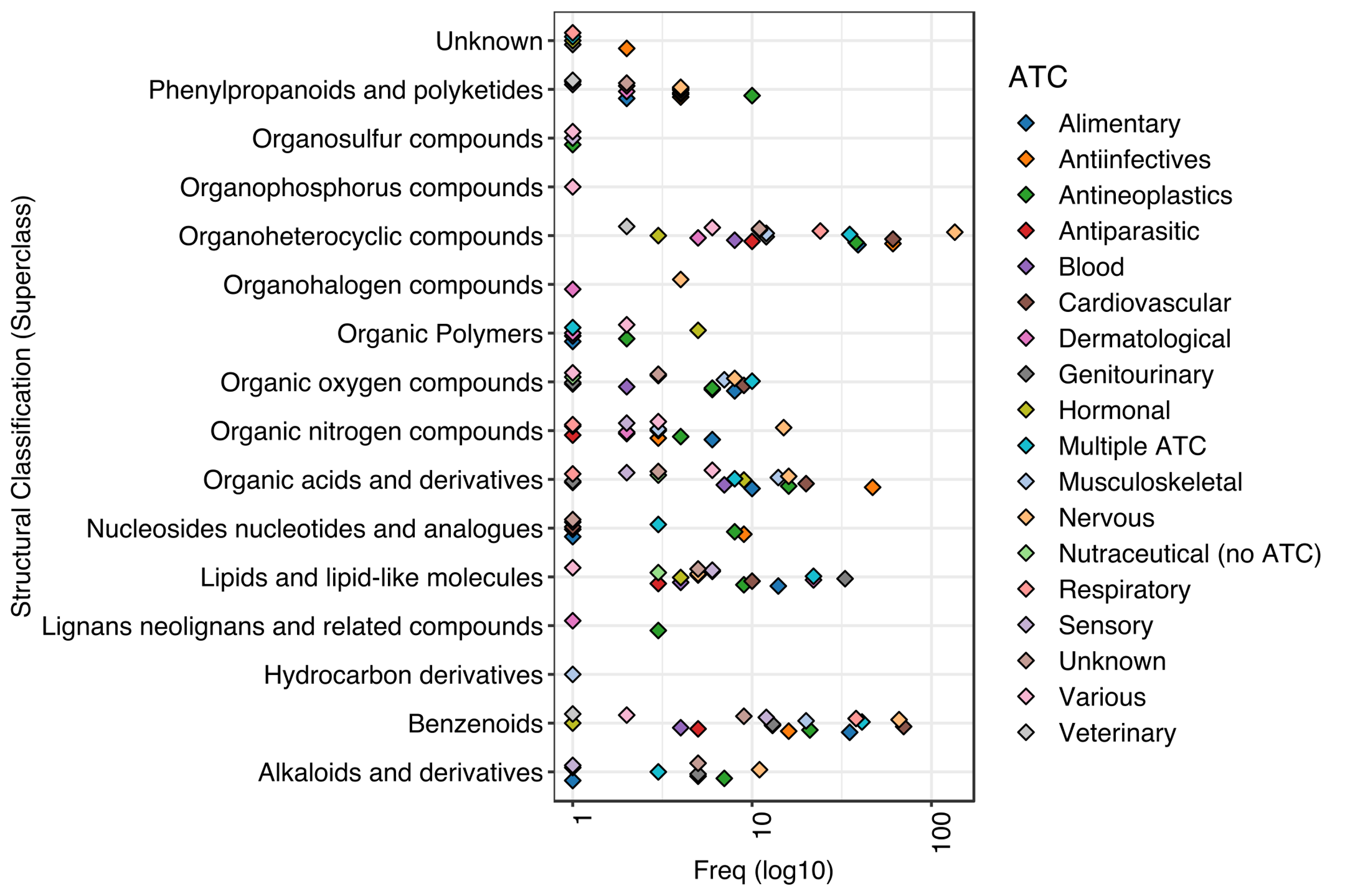

S1b

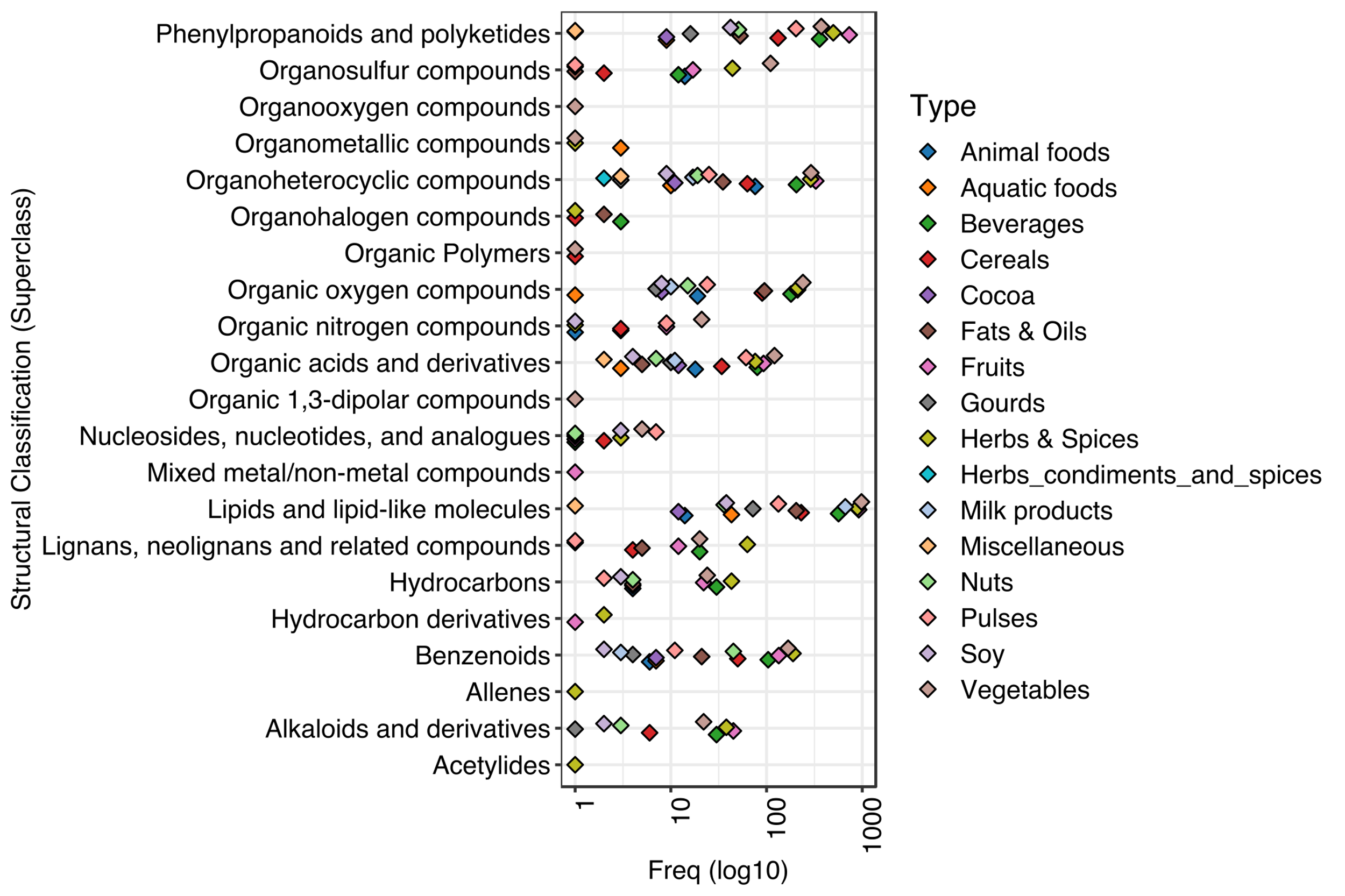

**Fig S2**: ‘Concordance at top’ (CAT plots)^55^ of fingerprints. We performed a rank-based assessment of drug-pairs called by each fingerprint as top scoring pairs and found a low concordance between them. The CAT plots here depict the concordance of each fingerprint (when taken as reference) against all other fingerprints. Each fingerprint was taken as a reference and the top 100 high scoring drug-pairs were ranked in decreasing order to estimate the overlapping proportions between each fingerprint when compared with the reference. Mathematically, for $ith$ top-ranked molecules, concordance is defined as $length(intersect\left( list1\left[ 1:i \right],list2\left[ 1:i \right] \right))/i$.

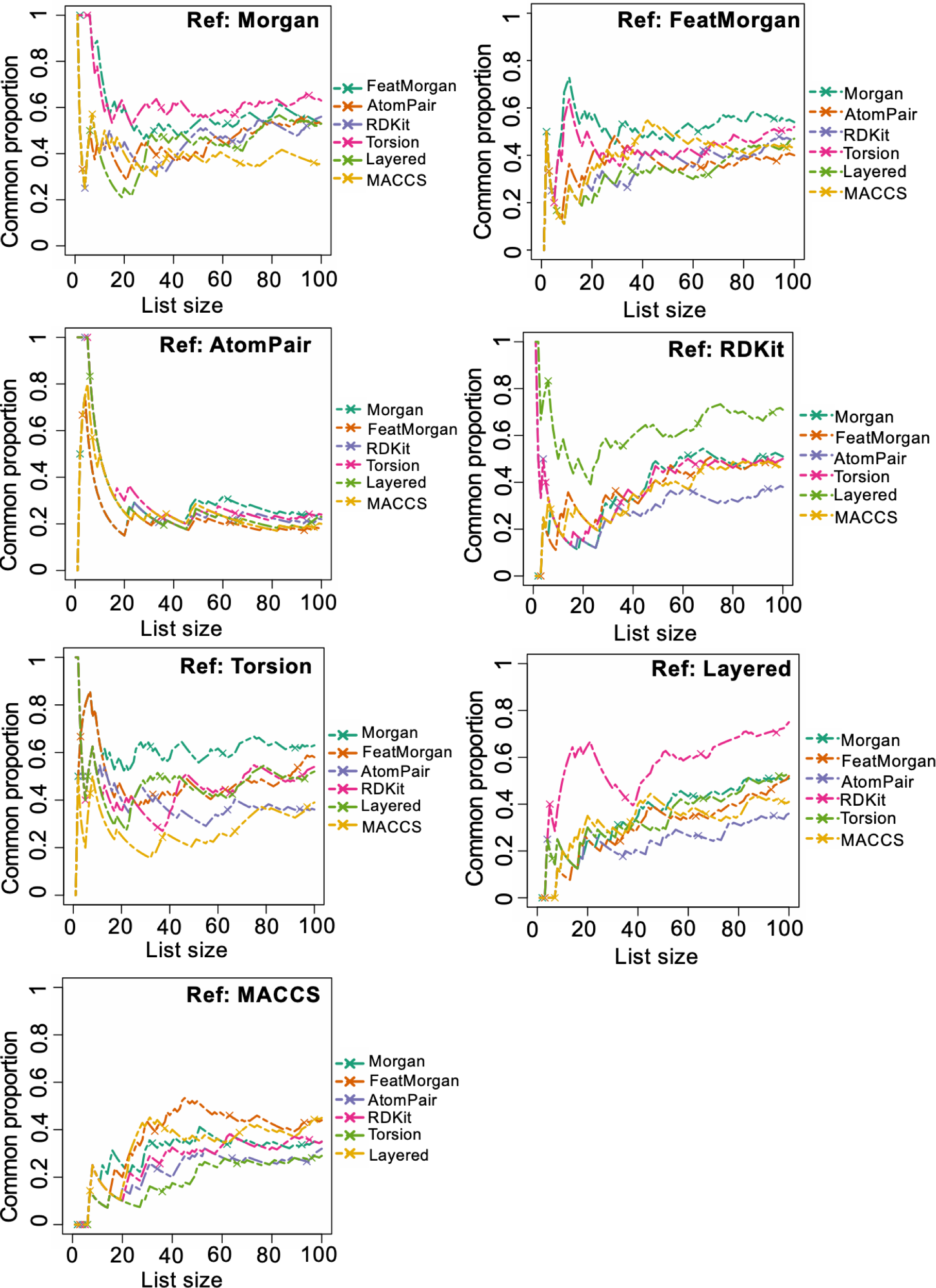

**Fig S3: Drug selection for training and test set split:** All drugs have superclass information (identified by their chemical class which they belong to). S3a plot has the counts of all drugs and their occurrence in each category. For creating the test set (completely independent from training set), step-1) we took top highly represented superclasses (at least > 100 drugs), 4 superclasses meet this threshold and in step-2) randomly selected 20% drugs from these 4 Superclasses (got 231 drugs). The number of these test set drugs in each superclass is represented in S3b.

S3a)

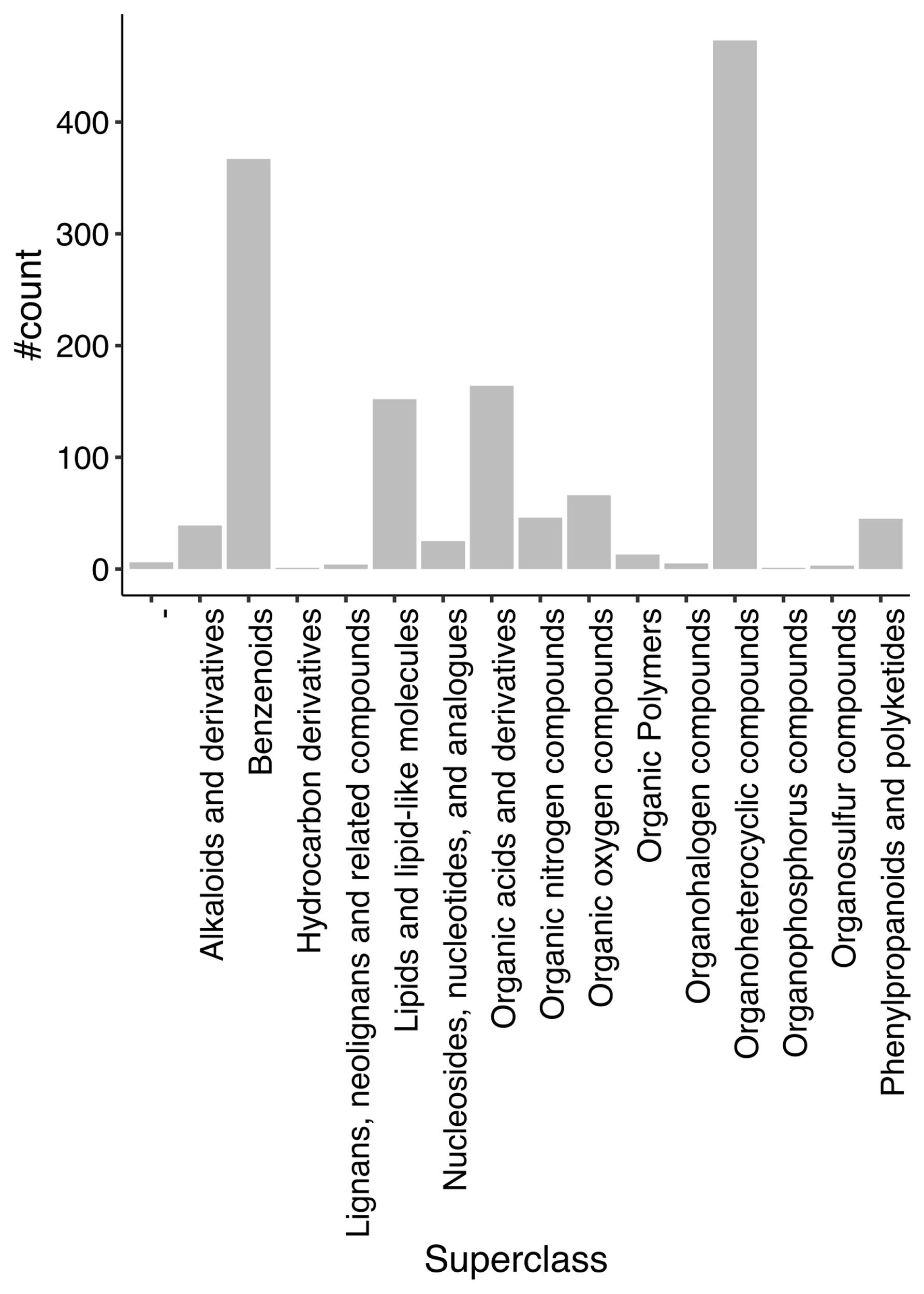

S3b) Numbers obtained after splitting - percentage of drugs (from the 4 superclasses) in the exclusive test set. Pairs of drugs were created in train set (1,179 drugs) and test set (231 drugs)

| **Superclass** | **Total count** | **In test set** |
| --- | --- | --- |
| Benzenoids | 367 | 66 (~18%) |
| Lipids and lipid-like molecules | 152 | 25 (~16%) |
| Organic acids and derivatives | 164 | 43 (~26%) |
| Organoheterocyclic compounds | 473 | 97 (~20%) |

**Fig S4: Random Forest hyperparameter tuning results:** hyperparameter tuning was performed with 10 iterations and 5-fold cross validation on training dataset. Four parameters were tuned number of trees (ntree), the number of observations at terminal nodes (*nodesize*), number of variables to split at each node (*mtry*) and class weights (*weight).* The runs were performed on cluster with parallel backend. Reported results are aggregated by ‘test.mean’ which takes the mean of the given performance measures over the number of cross validations for each iteration. The best performing hyperparameter combination was *ntree=241, nodesize=27, mtry=23, classwt=2515* which was used by the learner to train the final model.

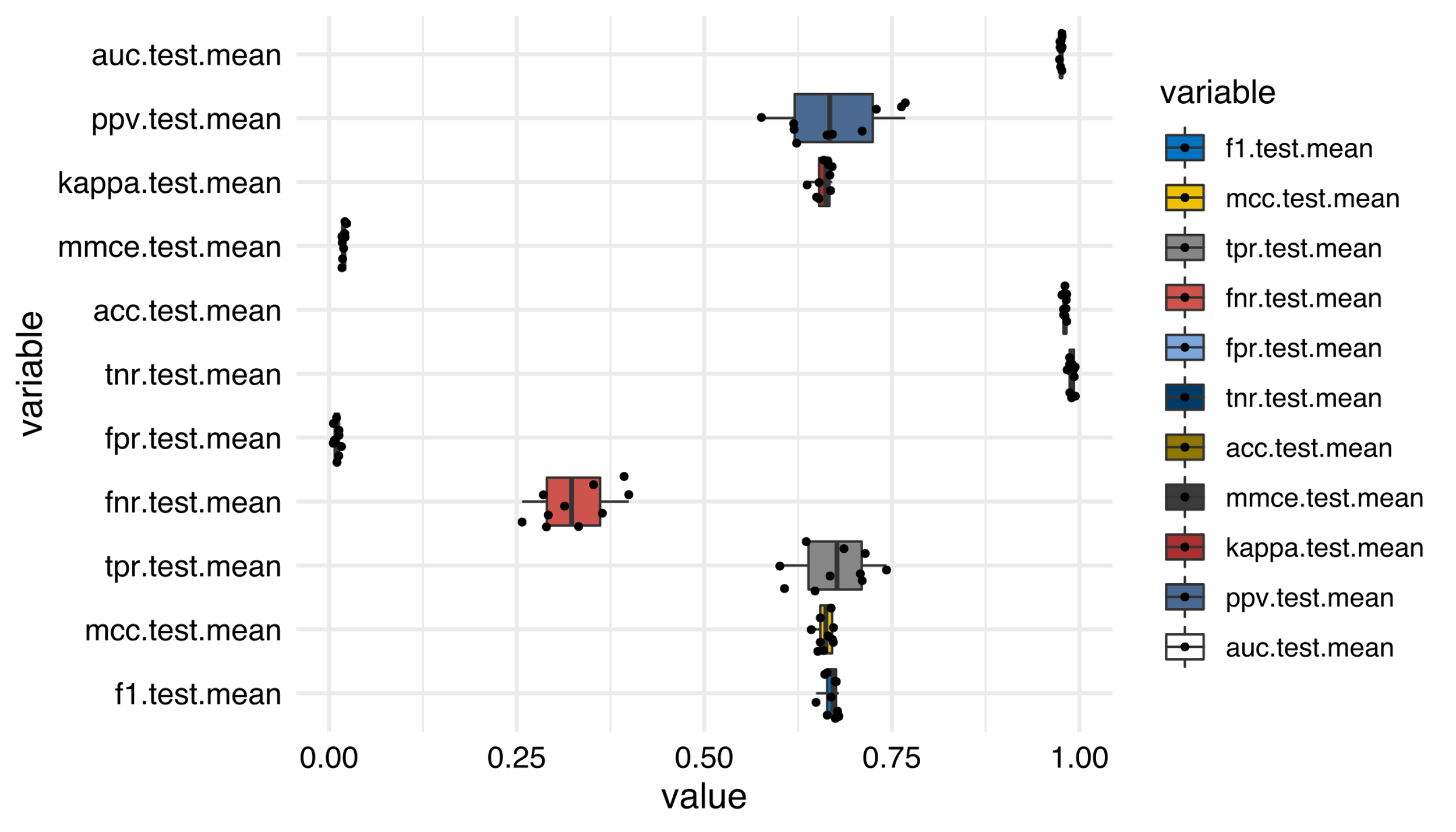

| Iter | ntree | nodesize | mtry | classwt | exec.time (sec) |
| --- | --- | --- | --- | --- | --- |
| 1 | 208 | 21 | 19 | 2358 | 190373.28 |
| 2 | 109 | 37 | 30 | 692 | 110327.98 |
| 3 | 106 | 47 | 29 | 1674 | 110495.07 |
| 4 | 109 | 49 | 15 | 2604 | 105582.19 |
| 5 | 103 | 43 | 21 | 1709 | 97638.94 |
| 6 | 229 | 21 | 21 | 1233 | 204165.72 |
| 7 | 241 | 27 | 23 | 2515 | 211519.6 |
| 8 | 199 | 37 | 22 | 2008 | 176792.84 |
| 9 | 298 | 31 | 27 | 1734 | 259921.88 |
| 10 | 279 | 48 | 30 | 1821 | 238525.75 |

**Fig S5** (A) Table describes the compounds tested in the cox-1 inhibitor assay, their source and usage. Triflusal is the drug which is known to bind to Cox-1 and it was a positive control in our experiment. The food compound which came out as hit with our prediction models is 5-methoxysalicylic acid (referred to as test compound) for which target engagement is being studied. Additional inclusion was 4-isopropylbenzoic acid (selected based on high FM score as compared to test compound but deemed no match by prediction models) as a negative control. (B) The reaction mechanism involves fluorometric detection of intermediate product (prostaglandin G2) generated by cox-enzyme.

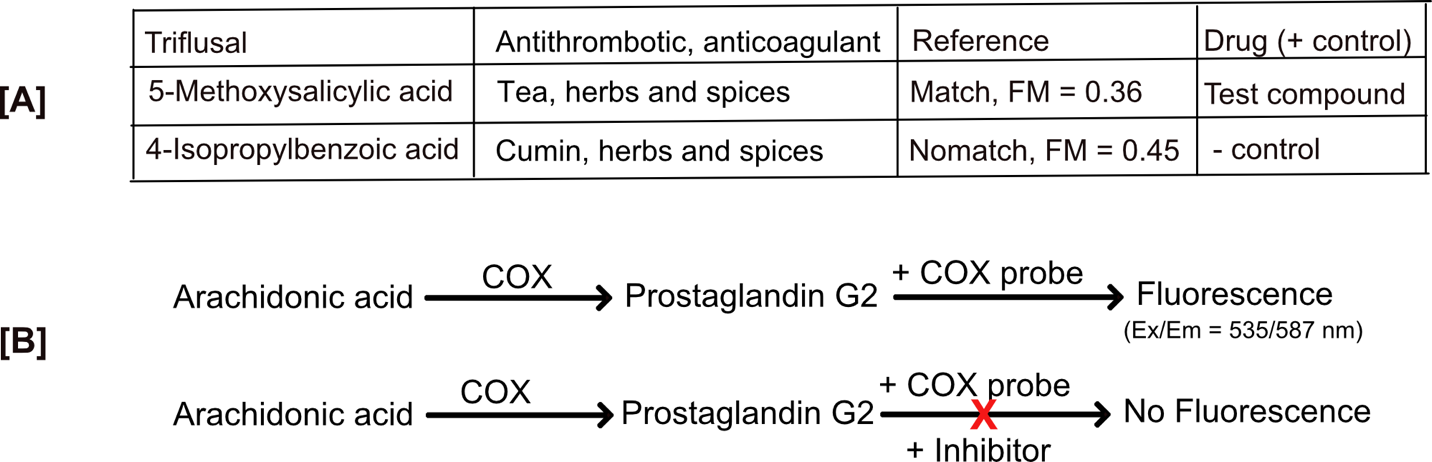

**Table S3 Model performance metrics and their description: TP – true positive, FP – false positive, TN – true negative, FN – false negative, TPR – true positive rate (TP/Positives), TNR – true negative rate (TN/negatives)**

| **Performance metric** | **Formula** |
| --- | --- |
| Matthews correlation coefficient (MCC) | $MCC= \frac{TPXTN-FPXFN}{\sqrt{\left( TP+FP \right)\left( TP+FN \right)(TN+FP)(TN+FN)}}$ |
| F1 score | $F1= \frac{2TP}{2TP+FP+FN}$ |
| Balanced accuracy (BAC) | $BAC= \frac{TPR+TNR}{2}$ |
| Kappa statistic | $K=\frac{2X(TPXTN-FNXFP)}{\left( TP+FP \right)X\left( FP+TN \right)+\left( TP+FN \right)X(FN+TN)}$ |
| Positive predictive value (PPV) | $PPV= \frac{TP}{TP+FP}$ |
| Accuracy | $ACC= \frac{TP+TN}{Positives+Negatives}$ |
| Mean misclassification error (MMCE) | Defined as: mean(response != truth) |

**Table S4** Cox-assay results. Relative Inhibition (%)

| Conc.(μM) | Conc. (log) | Triflusal | 5-Methoxysalicylic acid | 4-Isopropylbenzoic acid |
| --- | --- | --- | --- | --- |
| 400 | 2.60205999 | 50.2711928 | 44.5041162 | -106.98147 |
| 200 | 2.30103 | 53.7473155 | 41.4790048 | -38.982039 |
| 100 | 2 | 49.9543006 | 40.2151692 | -0.4554861 |
| 50 | 1.69897 | 44.7753213 | 36.7105663 | 10.6613523 |
| 25 | 1.39794001 | 41.6782677 | 32.5953934 | 14.4918464 |
| 12.5 | 1.09691001 | 41.3314119 | 33.0239321 | 17.6503106 |
